# Drift-diffusion models for multiple-alternative forced-choice decision making

**DOI:** 10.1101/542431

**Authors:** Alex Roxin

**Affiliations:** Barcelona Graduate School of Mathematics, Barcelona, Spain

**Keywords:** decision making, networks, winner-take-all

## Abstract

The canonical computational model for the cognitive process underlying two-alternative forced-choice decision making is the so-called drift-diffusion model (DDM). In this model, a decision variable keeps track of the integrated difference in sensory evidence for two competing alternatives. Here I extend the notion of a drift-diffusion process to multiple alternatives. The competition between *n* alternatives takes place in a linear subspace of *n –* 1 dimensions; that is, there are *n* – 1 decision variables, which are coupled through correlated noise sources. I derive the multiple-alternative DDM starting from a system of coupled, linear firing rate equations. If the original neuronal system is nonlinear, one can once again derive a model describing a lower dimensional diffusion process. The dynamics of the nonlinear DDM can be recast as the motion of a particle on a potential, the general form of which is given analytically for an arbitrary number of alternatives.

## Introduction

Perceptual decision making tasks require a subject to make a categorical decision based on noisy or ambiguous sensory evidence. A computationally advantageous strategy in doing so is to integrate the sensory evidence in time, thereby improving the signal-to-noise ratio. Indeed, when faced with two possible alternatives, accumulating the difference in evidence for the two alternatives until a fixed threshold is reached is an optimal strategy, in that it minimizes the mean reaction time for a desired level of performance. This is the computation carried out by the sequential probability ratio test devised by Wald [1], and its continuous-time variant, the drift-diffusion model (DDM) [2]. It would be hard to overstate the success of these models in fitting psychophysical data from both animals and human subjects in a wide array of tasks, e.g. [2, 3, 4, 5], suggesting that brain circuits can implement a computation analogous to the DDM.

At the same time, neuroscientists have characterized the neuronal activity in cortical areas of monkey, which appear to reflect an integration process during DM tasks [6]. The relevant computational building blocks, as revealed from decades of in-vivo electrophysiology, seem to be neurons, the activity of which selectively increases with increasing likelihood for a given upcoming choice. Attractor network models, built on this principle of competing, selective neuronal populations, generate realistic performance and reaction times; they also provide a neuronal description which captures some salient qualitative features of the in-vivo data [7, 8].

While focus in the neuroscience community has been almost exclusively on 2-alternative DM (although see [9]), from a computational perspective there does not seem to be any qualitative difference between 2 or more alternatives. In fact, increasing the number of alternatives is as trivial as adding another neuronal population to the competition. On the other hand, how to add an alternative to the DDM framework does not seem, on the face of things, obvious. Several groups have sought to link the DDM and attractor networks for 2-alternative DM. When the attractor network is assumed linear, one can easily derive an equation for a decision variable, representing the difference in the activities of the two competing populations, which precisely obeys a DDM [10]. Here I will illustrate how one can extend such a derivation to an arbitrary number of alternatives and thereby obtain an equivalent DDM. The relevant decision variables are combinations of the activity of the neuronal populations, and which represent distinct modes of competition.

Of course, linear models do not accurately describe the neuronal data from experiments on DM. However, previous work has shown that attractor network models for 2-alternative DM operate in the vicinity of pitchfork bifurcation, which is what underlies the winner-take-all competition leading to the decision dynamics [11]. In this regime the neuronal dynamics is well described by a stochastic normal form equation which right at the bifurcation is precisely equivalent to the DDM with an additional cubic nonlinearity. This *nonlinear* DDM fits behavioral data extremely well, including both correct and error reaction times. Here I will show how such normal form equations can be derived for an arbitrary number of neuronal populations. These equations can be thought of as nonlinear DDMs and, in fact, are identical to the linear DDMs with the addition of quadratic nonlinearities (for *n >* 2). Amazingly, the dynamics of such a nonlinear DDM can be recast as the diffusion of particle on a potential, which is obtained analytically, for arbitrary *n*.

## Results

The canonical drift diffusion model (DDM) can be written

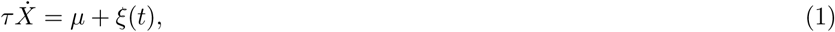

where *X* is the decision variable, and *µ* is the drift or the degree of evidence in favor of one choice over the other: we can associate choice 1 with positive values of *X* and choice 2 with negative values. The Gaussian process *ξ*(*t*) represents noise ′ and/or uncertainty in the integration process, with 〈*ξ*(*t*)〉 = 0 and 〈*ξ*(*t*)*ξ*(*t*′)〉 = *σ*^2^*δ*(*t* − *t*). I have also explicitly included a characteristic time-scale *τ*, which will appear naturally if one derives Eq.1 from neuronal model. The decision variable evolves until reaching one of two boundaries ±*θ* at which point a decision for the corresponding choice has been made.

It is clear that a single variable can easily be used to keep track of two competing processes by virtue of its having two possible signs. But what if there are three or more alternatives? In this case it is less clear. In fact, if we consider a drift-diffusion process such as the one in Eq.1 as an approximation to an actual integration process carried out by neuronal populations, then there is a principled approach to deriving the corresponding DDM. The value of such an approach is that one can directly tie the DDM to the neuronal dynamics, thereby linking behavior to neuronal activity.

I will first consider the derivation of a DDM starting from a system of linear firing rate equations. This analysis is similar to the that found in section 4.4 of [12], although the model of departure is different. In this case the derivation involves a rotation of the system so as to decouple the linear subspace for the competition between populations from the subspace which describes non-competitive dynamical modes. This rotation is equivalent to expressing the firing rates in terms of a set of orthogonal basis functions: one set for the competition, and another for the non-competitive modes. I subsequently consider a system of nonlinear firing rate equations. In this case one can once again derive a reduced set of equations to describe the decision-making dynamics. The reduced models have precisely the form of the corresponding DDM for a linear system, but now with additional nonlinear terms. These terms reflect the winner-take-all dynamics which emerge in nonlinear systems with multi-stability. Not only do the equations have a simple, closed-form solution for any number of alternatives, but they can be succinctly expressed in terms of a multivariate potential.

### Derivation of a DDM for 2-alternative DM

The DDM can be derived from a set of linear equations which model the competition between two populations of neurons, the activity of each of which encodes the accumulated evidence for the corresponding choice. The equations are

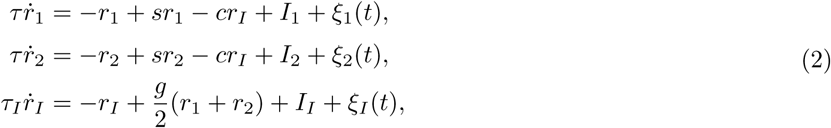

where *r*_*I*_ represents the activity of a population of inhibitory neurons. The parameter *s* represents the strength of excitatory self-coupling and *c* is the strength of the global inhibition. The characteristic time constants of excitation and inhibition are *τ* and *τ*_*I*_ respectively. A choice is made for 1 (2) whenever *r*_1_ = *r*_*th*_ (*r*_2_ = *r*_*th*_).

It is easier to work with the equations if they are written in matrix form, which is

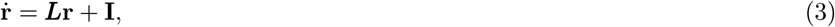

where

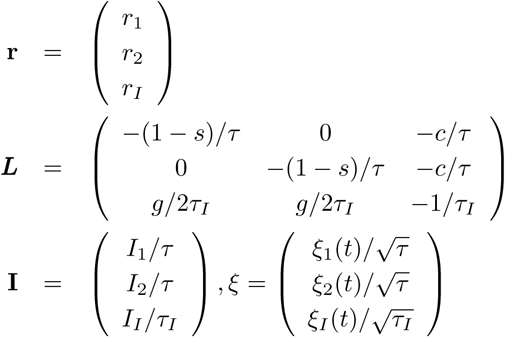

In order to derive the DDM, I express the firing rates in terms of three orthogonal basis functions: one which represents a competition between the populations e_1_, a second for common changes in population rates e_*c*_ and a third which captures changes in the activity of the inhibitory cells e_*I*_. Specifically we write

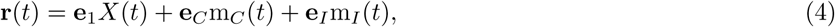

where e_1_ = (1, −1, 0) and e_C_ = (1, 1, 0) and e_I_ = (0, 0, 1). The decision variable will be *X*, while *m*_*C*_ and *m*_*I*_ stand for the common mode and inhibitory mode respectively.

The dynamics of each of these modes can be isolated in turn by projecting Eq.2 onto the appropriate eigenvector. For example, the dynamics for the decision variable *X* are found by projecting onto e_1_, namely

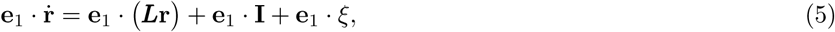

and similarly the dynamics for m_*c*_ and m_*I*_ and found by projecting onto e_*C*_ and e_*I*_ respectively. Doing so results in the set of equations

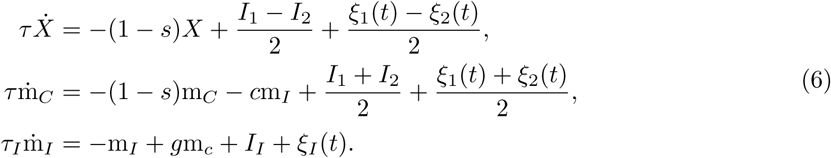

If the self-coupling is sufficiently strong, i.e. *s* = 1, then these equations simplify to

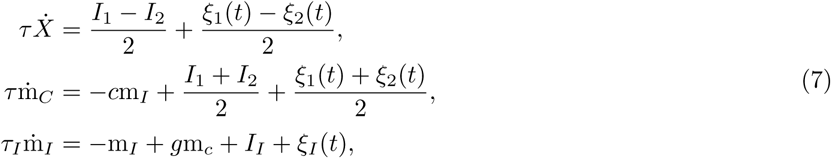

from which it is clear that the equation for *X* describes a drift-diffusion process. It is formally identical to Eq.1 with 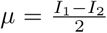 and 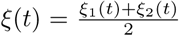. Importantly, *X* is uncoupled from the common and inhibitory modes, which themselves form a coupled subsystem. If one ignores the noise terms, the steady state of this linear system is (*X,* m_*C*_, m_*I*_) = (0, M_*c*_, M_*I*_) where

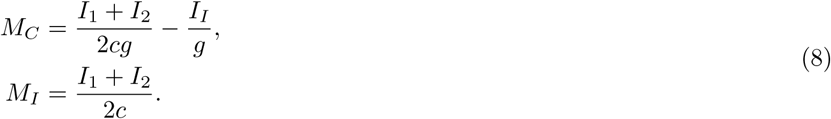

One can study the stability of this solution by considering a perturbation of the form (*X,* m_*C*_, m_*I*_) = (0, M_*c*_, M_*I*_) + (*δX, δ*m_*C*_ *, δ*m_*I*_)*e*^*λt*^. Plugging this into Eq.7 one finds that there is a zero-eigenvalue associated with the decision variable, i.e. *λ*_1_ = 0, whereas the eigenvalues corresponding to the subsystem comprising the common and inhibitory modes are given by

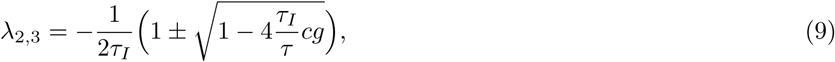

which always have a negative real part. Therefore, as long as *τ*_*I*_ is not too large, perturbations in the common mode or in the inhibition will quickly decay away. This allows us to ignore their dynamics and assume they take on their steady state values. Finally, the bounds for the decision variable are found by noting that *r*_1_ = *X* + M_*C*_ and *r*_2_ = −*X* + M_*C*_. Therefore, given that the neuronal threshold for a decision is defined as *r*_*th*_, we find that *θ* = ±(*r*_*th*_ − M_*C*_).

### Derivation of a DDM for 3-alternative DM

I will go over the derivation of a drift-diffusion process for 3-choice DM in some detail for clarity, although conceptually it is very similar to the 2-choice case. Then the derivation can be trivially extended to n-alternative DM for any n.

The linear rate equations are once again given by Eq.3, with

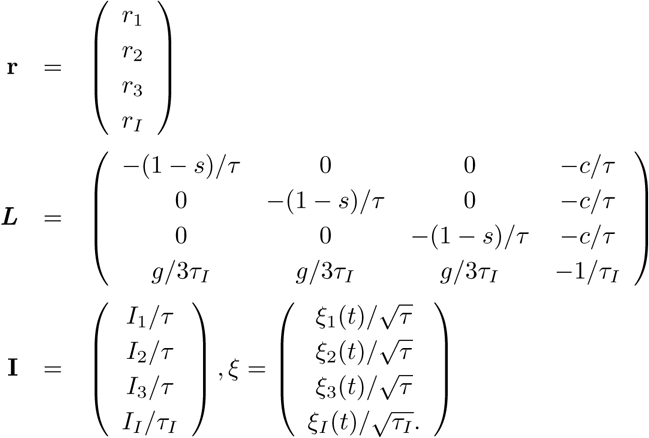

One once again writes the firing rates in terms of orthogonal basis functions, of which there must now be four. The common and inhibitory modes are the same as before, whereas now there will be two distinct modes to describe the competition between the three populations, in contrast to just a single decision variable. Specifically **r** = e_1_*X*_1_(*t*) + e_2_*X*_2_(*t*) + e_*C*_ m_*C*_(*t*) + e_*I*_ m_*I*_, where

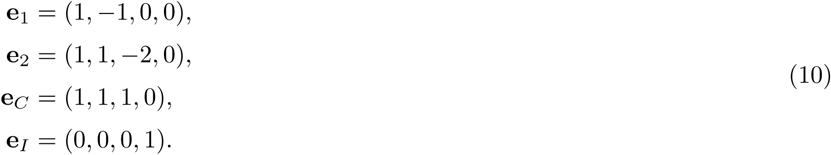

One projects Eq.3 onto the four relevant eigenvectors, which leads to the following equations

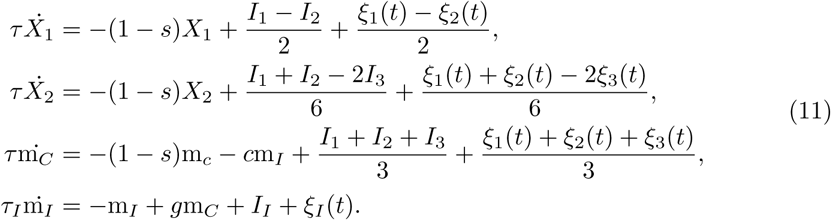

When *s* = 1 then the first two equations in Eq.11 describe a drift-diffusion process in a two-dimensional subspace, while the coupled dynamics of the common and inhibitory modes are once again strongly damped. The DDM for 3-alternative DM can therefore be written

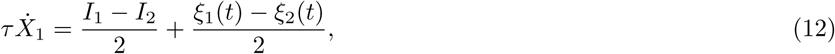

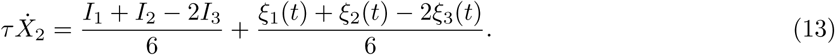

Note that the dynamics of the two decision variables *X*_1_ and *X*_2_ are coupled through the correlation in their noise sources. The decision boundaries are set by noting that

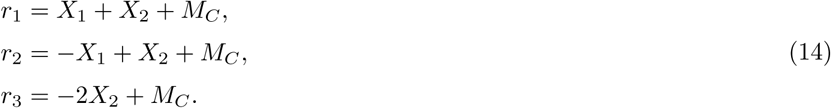

Therefore, given that the neuronal threshold for a decision is defined as *r*_*th*_ we can set three decision boundaries: 1 - Population 1 wins if *X*_2_ = −*X*_1_ + *r*_*th*_ − *M*_*C*_, 2 - Population 2 wins if *X*_2_ = *X*_1_ + *r*_*th*_ − *M*_*C*_ and 3 - Population 3 wins if *X*_2_ = −(*r*_*th*_ − *M*_*C*_)/2. These three boundaries define a triangle in (X,Y)-space over which the drift-diffusion process take place.

### Derivation of DDMs for n-alternative DM

The structure of the linear rate equations Eq.3 can be trivially extended to any number of competing populations. In order to derive the corresponding DDM we need only properly define the basis functions for the firing rates. The common and inhibitory modes always have the same structure. Therefore, we must find the *n* − 1 mutually-orthogonal basis functions for the competition process itself. These eigenfunctions always have the property that the sum of the elements is equal to zero. If we call these basis functions e_i_ and the corresponding decision variables *X*_*i*_ for *i* = [1*, n* − 1], then the firing rates are 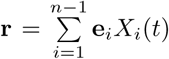 and it is easy to show that the dynamics for the *k*-th decision variable are given by

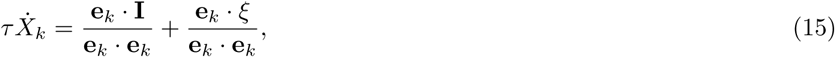

as long as *s* = 1. The decision boundaries are defined by setting the firing rates equal to their threshold value for a decision, i.e.

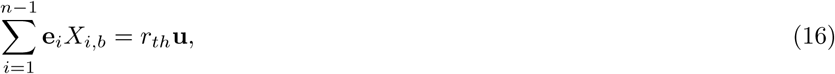

where **u** = (1, 1*, …,* 1).

One basis set for *n*-alternative DM, is to take for the k-th eigenvector

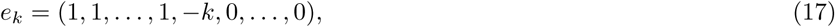

where the element −*k* appears in the (*k* + 1)-st spot, and which is a generalization of the eigenvector basis taken earlier for 2- and 3-alternative DM. With this choice, the equation for the *k*-th decision variable can be written

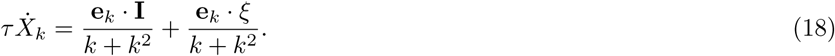

The firing rate for the *i*th *neuronal* population can then be expressed in terms of the decision variables as

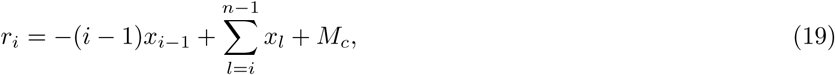

which, given a fixed neuronal threshold *r*_*th*_, directly gives the bounds on the decision variables. Namely, the *i*-th neuronal population wins when 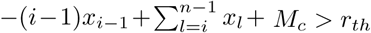.

The n-alternative DDM reproduces the well-known Hick’s law [13], which postulates that the mean reaction time (RT) increases as the logarithm of the number of alternatives, for fixed performance, see Fig.1.

**Figure 1.**
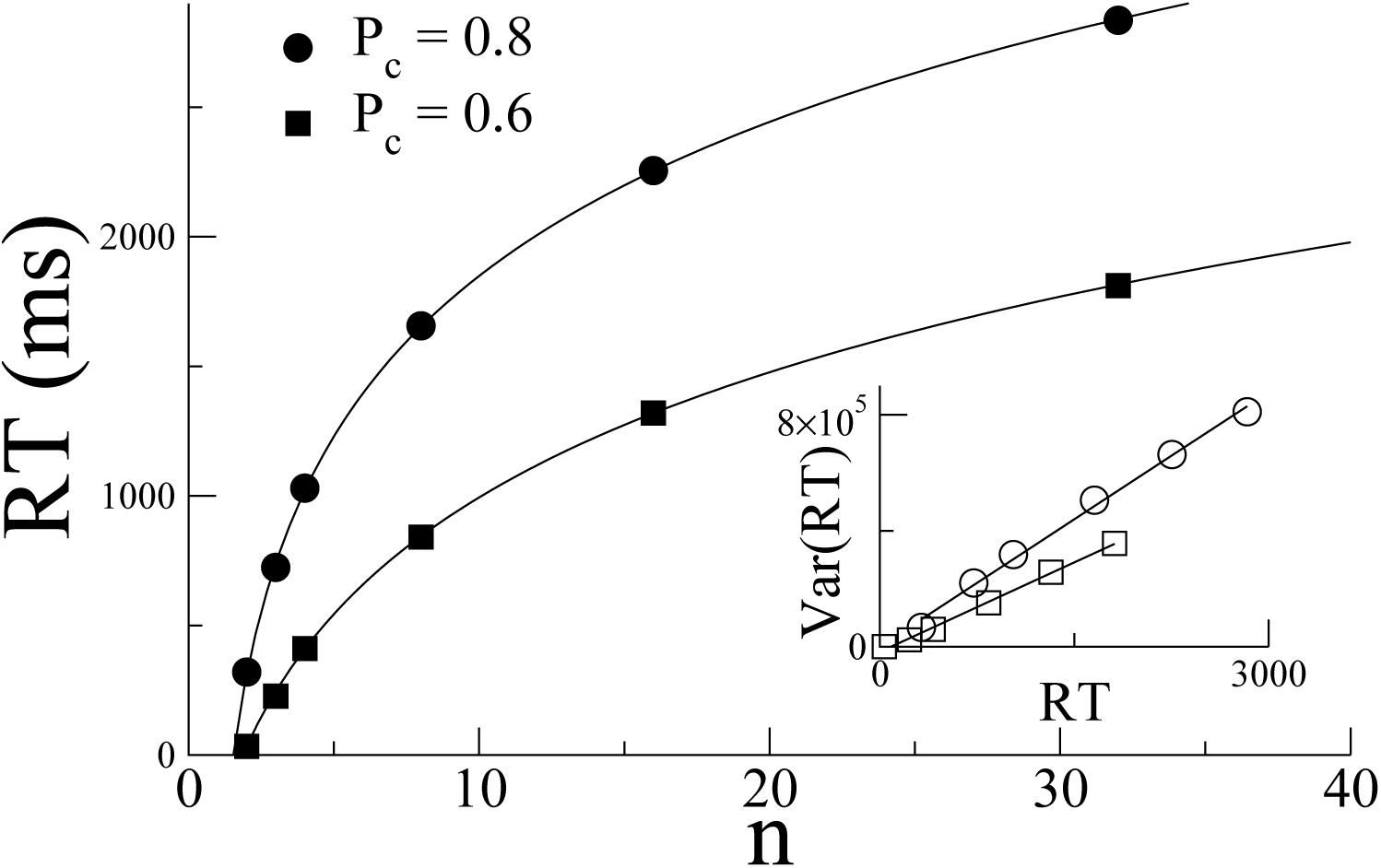
Hick’s Law for multi-alternative DDMs. If the performance is held fixed as the number of alternatives *n* is increased, then the mean reaction time increases as the logarithm of *n.* This is shown for both a performance of 0.8 (circles) as well as 0.6 (squares). The inset shows that the variance in the RT increases proportional to the mean RT. Parameter values are σ = 1, τ = 20ms. The thresholds *θp*_*c*_=0.8 = 3 and *θP_*c*_=0*.6 = 0.9 for *n* = 2, and are increased for *n* > 2 to achieve the same performance. The initial condition is always taken to be *X_*i*_ =* 0 for all i. The symbols are the average of 10,000 runs. The solid curves in the main panel are fits to the function *RT = a* + *b* ln (*c* + *n*). The solid lines in the inset are best-fit linear regressions.

## Derivation of a reduced model for 2-choice DM for a nonlinear system

A more realistic firing rate model for a decision-making circuit allows for a nonlinear input-output relationship in neuronal activity. For 2-alternative DM the equations are

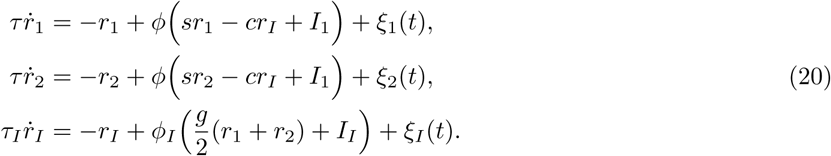

The nonlinear transfer function *φ* (*φ*_*I*_) does not need to be specified in the derivation. The noise sources *ξ*_*i*_ are taken to be Gaussian white noise and hence must sit outside of the transfer function; they therefore directly model fluctuations in the population firing rate rather than input fluctuations. Input fluctuations can be modelled by allowing for a non-white noise process and including it directly as an additional term in the argument of the transfer function. Again, this would not affect the derivation itself.

The details of the derivation for two alternatives can be found in [11], but here I give a flavor for how one proceeds; the process will be similar when there are three or more choices. One begins by ignoring the noise sources and linearizing Eqs.20 about a fixed point value for which the competing populations have the same level of activity, and hence also *I*_1_ = *I*_2_. Specifically one takes (*r*_1_*, r*_2_*, r*_I_) = (*R, R, R*_*I*_) + (*δr*_1_*, δr*_2_*, δr*_*I*_)*e*^*λt*^, where *δr* ≪ 1. In vector form this can be written **r** = **R** + *δ***r***e*^*λt*^. Plugging this ansatz into Eqs.20 and keeping only terms linear in the perturbations leads to the following system of linear equations

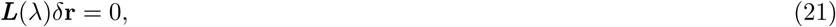

where

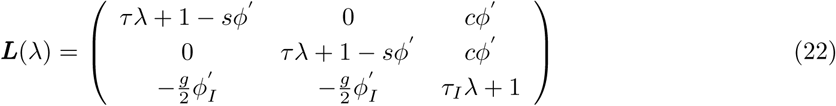

and the slope of the transfer function *φ*′ is calculated at the fixed point. Note that the matrix Eq.22 has a very similar structure to the linear operator in Eq.3. This system of equations only has a solution if the determinant of the matrix is equal to zero; this yields the characteristic equation for the eigenvalues *λ*. These eigenvalues are

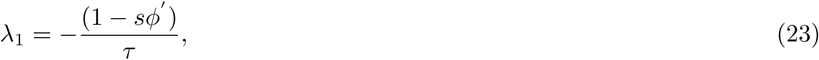

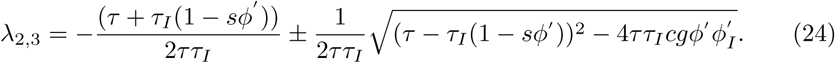

Note that *λ*_1_ = 0 if 1 − *sφ*′ = 0, while the real part of the other two eigenvalues is always negative. This indicates that there is an instability of the fixed point in which the activity of the neuronal populations is the same, and that the direction of this instability can be found by setting 1 − *sφ*′ = 0 and *λ* = 0 in Eq.22. This yields ***L***_0_ ⋅ *δ***r** = 0, where

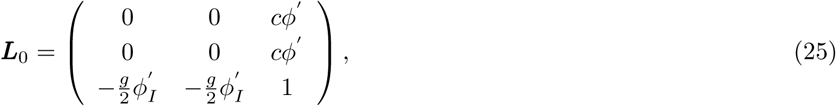

the solution of which can clearly be written *δ***r** = (1, −1, 0). This is the same competition mode found earlier for the linear system.

### A brief overview of normal-form derivation

At this point it is still unclear how one can leverage this linear analysis to derive a DDM. Specifically, and unlike in the linear case, one cannot simply rotate the system to uncouple the competition dynamics from the non-competitive modes. Also, note that the steady states in a nonlinear system depend on the external inputs, whereas that is not the case in a linear system. In particular, the DDM has a drift term *µ* which ought to be proportional to the difference in inputs *I*_1_ − *I*_2_, whereas to perform the linear stability we assumed *I*_1_ = *I*_2_. Indeed, if we assume the inputs are different then the fixed-point structure is completely different. The solution is to assume that the inputs are only slightly different, and formalize this by introducing the small parameter *ϵ*. Specifically, we write 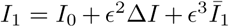, and 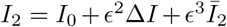, where ∆*I* is a change in the mean input to both populations. The firing rates are then also expanded in orders of *ϵ* and written

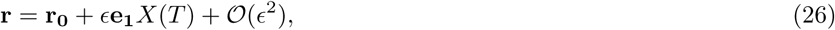

where r_0_ are the fixed point values, e_1_ is the eigenvector corresponding to the zero-eigenvalue and *X* is the decision variable which evolves on a slow time scale *T* = *ϵ*^2^*t*. The slow time scale arises from the fact that there is an eigenvector with zero eigenvalue; when we change the parameter values slightly, proportional to *ϵ*, the growth rate of the dynamics along that eigenvector is no longer zero, but still very small, in fact proportional to *ϵ*^2^ in this case.

The method for deriving the normal form equation, i.e. the evolution equation for *X*, involves expanding Eqs.20 in *ϵ*. At each order in *ϵ* there is a set of equations to be solved; at some orders, in this case first at order 𝓞(*ϵ*^3^), the equations cannot be solved and a solvability condition must be satisfied, which leads to the normal form equation.

### The normal-form equation for 2-choices

Following the methodology described in the preceding section leads to the evolution equation for the decision variable *X*

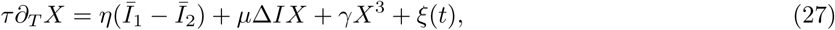

where for the case of Eqs.20, *η* = *φ′ /*2, *ξ*(*t*) = (*ξ*_1_(*t*) − *ξ*_2_(*t*))*/*2 and the coefficients *µ* and *γ* are

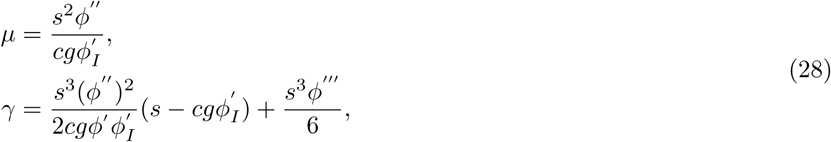

see [11] for a detailed calculation. Eq.27 provides excellent fits to performance and reaction-times for monkeys and human subjects, see Fig.3 from [11].

It is important to note that the form of Eq.27 only depends on there being a two-way competition, not on the exact form of the original system. As an example, consider another set of firing rate equations

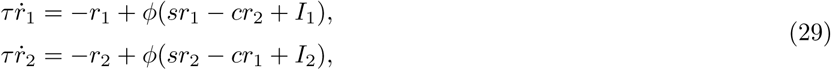

where rather than model the inhibition explicitly, an effective inhibitory interaction between the two populations is assumed. In this case the resulting normal-form equation is still Eq.27. In fact, performing a linear stability analysis on Eqs.29 yields a null eigenvector *e*_1_ = (1, −1). This indicates that the instability causes one population to grow at the expense of the other, in a symmetric fashion, as before. This is the key point which leads to the normal-form equation. More specifically we see that for both systems *r*_1_ = *R* + *X* while *r*_2_ = *R* − *X*, which means that if we flip the sign on the decision variable *X* and switch the labels on the neuronal populations, the dynamics are once again the same. This reflection symmetry ensures that all terms in *X* will have odd powers in Eq.27 [11]. It is broken only when the inputs to the two populations are different, i.e. by the first term on the r.h.s. in Eq.27.

As we shall see, the stochastic normal-form equation, Eq.27, which from now on I will refer to as a nonlinear DDM, has a very different form from the nonlinear DDMs for *n >* 2. The reason is, again, the reflection symmetry in the competition subspace for *n* = 2, which is not present for *n >* 2. Therefore, for *n >* 2 the leading-order nonlinearity is quadratic, and, in fact, a much simpler function of the original neuronal parameters.

## Three-alternative forced-choice decision making

The derivation of the normal form, and the corresponding DDM for three-choice DM differs from that for two-alternative DM in several technical details; these differences continue to hold for n-alternative DM for all *n* ≥ 3. Therefore I will go through the derivation in some detail here and will then extend it straightforwardly to the other cases.

Again we will make use of a particular system of firing rate equations to illustrate the derivation. We take a simple extension of the firing rate equations for two-alternative DM Eqs.20. The equations are

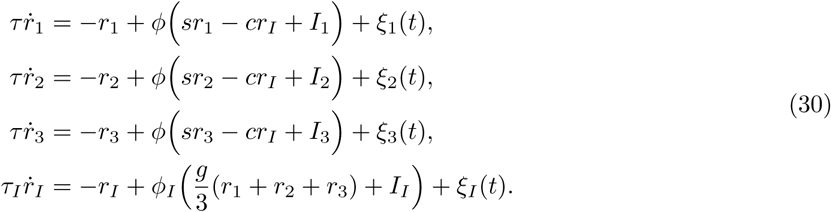

We first ignore the noise terms and consider the linear stability of perturbations of the state in which all three populations have the same level of activity (and so *I*_1_ = *I*_2_ = *I*_3_ = *I*_0_), i.e. **r** = **R** + *δ*r*e*^*λt*^, where **R** = (*R, R, R, R*_*I*_). This once again leads to a set of linear equations ***L***(*λ*)*δ***r** = 0. The 4th order characteristic equation leads to precisely the same eigenvalues as in the two-choice case, Eqs.23-24 with the notable difference that the first eigenvalue has multiplicity two. This means that if 1−*sφ′* = 0 then there will be two eigenvalues identically equal to zero and two stable eigenvalues. This is the first indication that the integration process underlying the DM process for three-choices will be two-dimensional. The eigenvectors for the DM process are found by solving ***L***_0_*δ***r** = 0, where

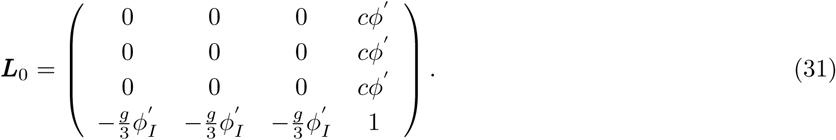

There are many possible solutions; a simple choice would be e_1_ = (1, −1, 0, 0) and e_2_ = (1, 1, −2, 0), and so 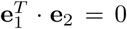. Note that in this linear subspace any valid choice of eigenvector will have the property that the sum of all the elements will equal zero; this will be true whatever the dimensionality of the DM process and reflects the fact that all of the excitatory populations excite the inhibitory interneurons in equal measure.

To derive the normal form I once again assume that the external inputs to the three populations differ by a small amount, namely 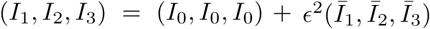, and then expand the firing rates as 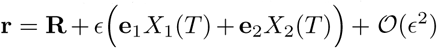, where the slow time *T* = *ϵt*. We expand Eqs.30 in orders of *ϵ*. In this case a solvability condition first arises at order *ϵ*^2^, which also accounts for the different scaling of the slow-time compared to 2-choice DM. Note that there are two solvability conditions, corresponding to eliminating the projection of terms at that order onto both of the left-null eigenvectors of ***L***_0_. As before, the left-null eigenvectors are identical to the right-null eigenvectors. Applying the solvability condition yields the normal form equations

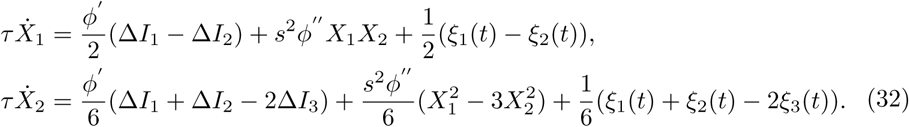

The nonlinear DDM Eq.32 provides an asymptotically correct description of the full dynamics in Eqs.30 in the vicinity of the bifurcation leading to the decision-making behavior. Fig.2A shows a comparison of the firing rate dynamics with the nonlinear DDM right at the bifurcation. The appropriate combinations of the two decision variables *X*_1_ and *X*_2_ clearly track the rate dynamics accurately, including the correct choice (here population 2) and reaction time. The nonlinear drift-diffusion process evolves in a triangular section of the plane, see Fig.2B.

**Figure 2.**
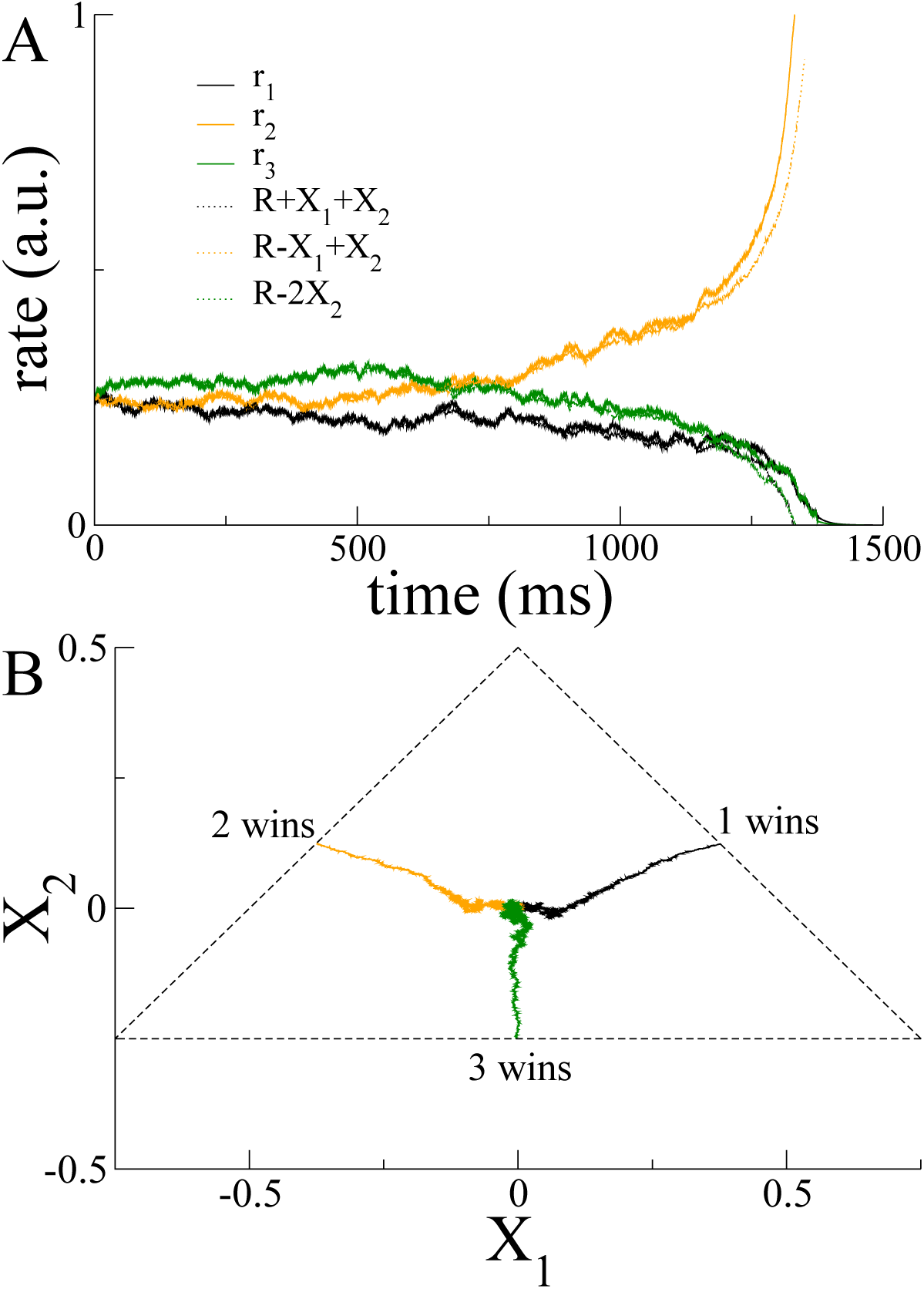
Decision-making dynamics for 3-alternatives A. **A** comparison of the dynamics from integration of the coupled firing rate equations Eqs.30 (solid lines) and from the DDM, Eq.32 (dotted lines). B. The dynamics from three separate simulations of the DDM, Eq.32, indicating three different winning alternatives. Parameters are: τ = 20ms, τ_*I*_ = 10ms, *s* = *c* = *g* =1, σ *=* 0.01. The function φ(*x*) *=* 0 for *x* < 0, *x^2^* for 0 ≤ *x* ≤ 1 and 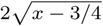 for *x* > 1. For this choice of transfer function and these parameter values, there is a bifurcation for *I*_0_ = 1/2, which is the value taken here. The steady-state firing rate *R =* 1/4. The threshold *θ* = 1/2.

### A note on the difference between the nonlinear DDM for 2A and 3A DM

The dynamics of the nonlinear DDM for 2A, Eq.27, depends strongly on the sign of the cubic coefficient *γ*. Specifically, when *γ <* 0 the bifurcation is supercritical, while for *γ >* 0 it is subcritical, indicating the existence of a region of multistability for ∆*I <* 0. In fact, in experiment, cells in parietal cortex which exhibit ramping activity during perceptual DM tasks, also readily show delay activity in anticipation of the sacade to their response field [14]. One possible mechanism for this would be precisely this type of multistability. When ∆*I* = 0, i.e. when the system sits squarely at the bifurcation, Eq.27 is identical to its linear counterpart with the sole exception of the cubic term. For *γ <* 0 the state *X* = 0 is stabilized. In fact, the dynamics of the decision variable can be viewed as the motion of a particle on a potential, which for *γ <* 0, increases rapidly as X grows, pushing the particle back. On the other hand, for *γ >* 0 the potential accelerates the motion of *X*, pushing it off to ±∞. This is very similar to the potential for the linear DDM with absorbing boundaries. Therefore, the nonlinear DDM for two-alternatives is qualitatively similar to the linear DDM when it is subcritical, and hence when the original neuronal system is multistable.

On the other hand, the nonlinear DDM for three-alternatives, Eq.32, has a much simpler, quadratic nonlinearity, the sign of which is always positive ^[1]^. The consequence of this is that there are no stable fixed points and the decision variables always evolve to ±∞. Furthermore, to leading order there is no dependence on the mean input, indicating that the dynamics are dominated by the behavior right at the bifurcation ^[2]^. The upshot is that Eq.32 is as similar to the corresponding linear DDM with absorbing boundaries as possible for a nonlinear system without fine tuning. This remains true for all *n >* 2.

This also means that neuronal systems with inhibition-mediated winner-take-all dynamics are generically multistable for *n >* 2, although for *n* = 2 they need not be. This is due to the reflection symmetry present only for *n* = 2.

## n-alternative forced-choice decision making

We can now extend the analysis for 3-alternative DM to the more general n-choice case. Again I start with a set of firing rate equations

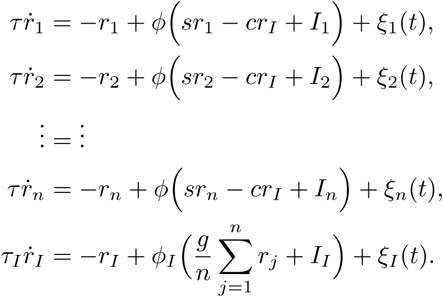

A linear stability analysis shows that the eigenvalues of perturbations of the state r = (*R, R, …, R, R*_I_) are given by Eqs.23 and 24, where the first eigenvalue has multiplicity *n* − 1. Therefore the decision making dynamics evolve on a manifold of dimension *n* − 1. The linear subspace associated with this manifold is spanned by *n* − 1 eigenvectors which are mutually orthogonal and the elements of which sum to zero. For *n*-alternatives, we take *n* − 1 eigenvectors of the form *e*_k_ = (1, 1*, …,* 1, −*k,* 0*, …,* 0), for the *k*-th eigenvector (again, the −*k* sits in the (k+1)-st spot). Therefore we can write

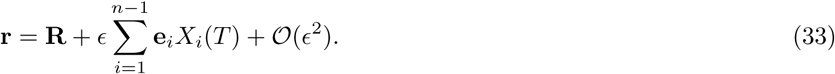

Following the same procedure as in the case of 3-choice DM ad applying the *n* − 1 solvability conditions at order 𝓞(*ϵ*^2^), one arrives at the following normal-form equation for the *k*-th decision variable for *n*-alternatives

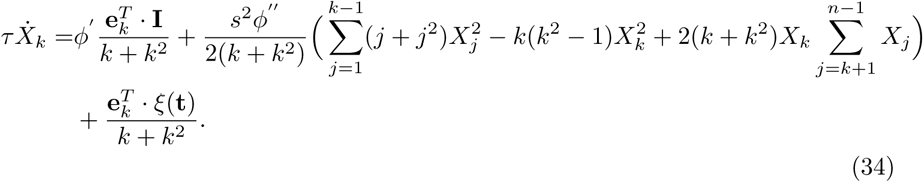

It is clarifying to write this formula out explicitly for some of the decision variables.

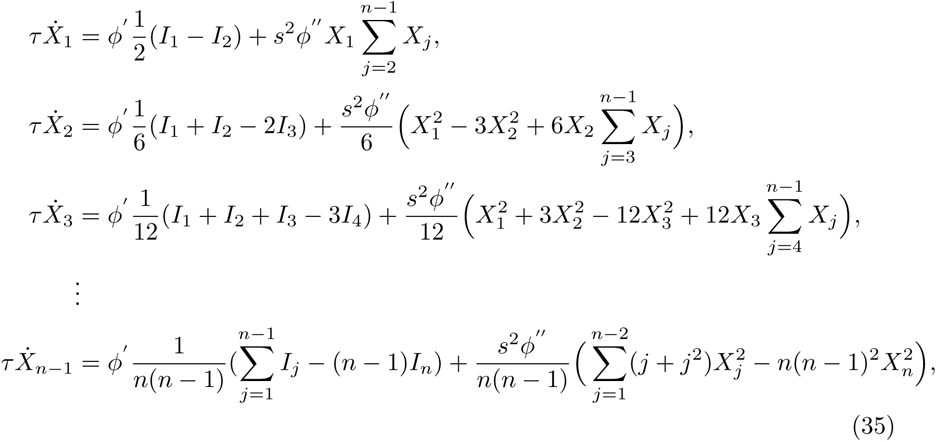

where I have left off the noise terms for simplicity.

Surprisingly, the *n* − 1 equations for the decision variables in *n*-alternative DM can all be derived from a single, multivariate function

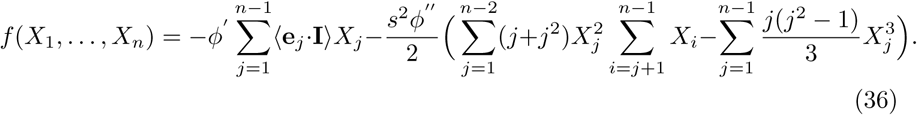

Then the equation for the *k*-th decision variable is simply

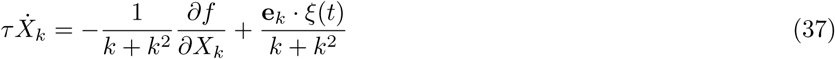

The dynamics of the function *f* is given by

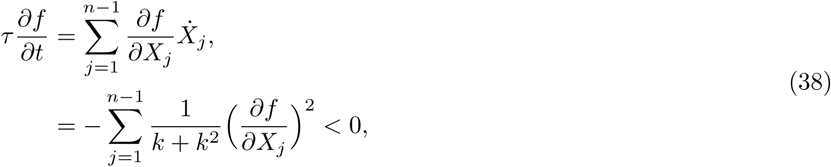

where I have ignored the effect of noise. Therefore the dynamics of the decision variables can be thought of as the motion of a particle on a potential landscape, given by *f*. Noise sources lead to a diffusion along this landscape.

## Discussion

In this paper I have illustrated how to derive drift-diffusion models starting from models of neuronal competition for n-alternative decision-making tasks. In the case of linear systems, the derivation consists of nothing more than a rotation of the dynamics onto a subspace of competition modes. This idea is not new, e.g. [12], although I have made the derivation explicit here, and have chosen as a model of departure one in which inhibition is explicitly included as a dynamical variable. For nonlinear systems, the corresponding DDM is a stochastic normal form, which is obtained here using the method of multiple-scales [15]. The nonlinear DDM was obtained earlier for the special case of two-alternative DM [11]. For four-alternative DM the nonlinear DDM was obtained with a different set of competitive basis functions [16] to describe performance and reaction time from experiments with human subjects. This led to a different set of coupled normal form equations from those given by Eq.34, although the resulting dynamics are, of course, the same. The advantage of the choice I have made in this paper for the basis functions, is that they are easily generalizable for any *n*, leading to a simple, closed-form expression for the nonlinear DDM for any arbitrary number of alternatives, Eq.34.

An alternative approach to describing behavior in DM tasks, is to develop a statistical or probabilistic description of evidence accumulation, e.g. [12, 17]. Such an approach also often leads to a drift-diffusion process in some limit. From a neuroscience perspective, however, it is of interest to pin down how the dynamics of neuronal circuits might give rise to animal behavior which is well described by a drift-diffusion process. This necessitates the analysis of neuronal models at some level. What I have shown here is that the dynamics in a network of *n* neuronal populations which compete via a global pool of inhibitory interneurons, can in general be formally reduced to a nonlinear DDM of dimension *n* − 1. The nonlinear DDMs differ from the linear DDMs through the presence of a single quadratic nonlinearity which accelerates the winner-take-all competition. In practical terms this nonlinear acceleration serves that same role as the hard threshold in the linear DDMs. Therefore the two classes of DDMs have quite similar behavior.

The DDMs derived here, both linear and nonlinear, are simple in form and should prove useful both for running Monte-Carlo simulations, as well as for formal mathematical analysis.

## Appendix

### Derivation of the normal form for 3-alternative DM

I assume the inputs to the three competing neuronal populations in Eqs.30 are slightly different, namely 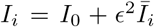 for *i* = {1, 2, 3}, which defines the small parameter *ϵ* ≪ 1. The firing rates are expanded as **r** = **R** + *ϵ***r**_1_ + *ϵ*^2^**r**_2_ + 𝓞(*ϵ*^3^) where **R** are the values of the rates at the fixed point. Finally, we define a slow time *T* = *ϵt*. Plugging the expansions for the inputs and rates into Eqs.30 and expanding leads to the following system of equations

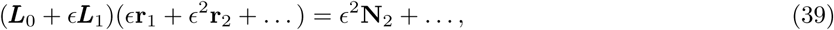

where to place the system at the bifurcation to winner-take-all behavior I choose 1 − *sφ′* = 0. Then the linear operator

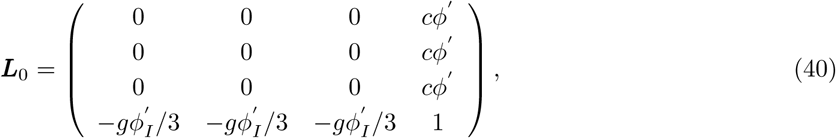

and as well

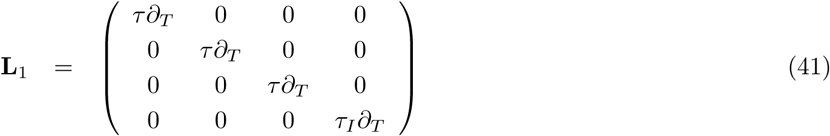

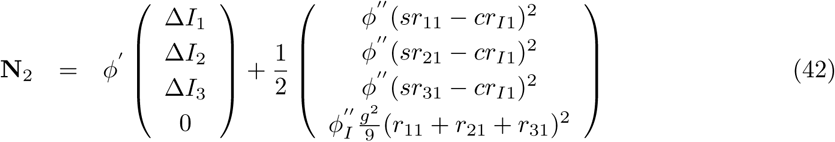

Now one solves the equations order by order.

𝓞(*ϵ*) :

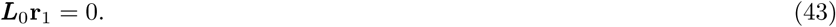

The solution can be written **r**_1_ = e_1_*X*_1_(*T*) + e_2_*X*_2_(*T*), where the right-null eigen-vectors of ***L***_0_ are e_1_ = (1, −1, 0, 0) and e_2_ = (1, 1, −2, 0). The left-null eigenvectors, which satisfy 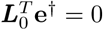 are identical to the right-null eigenvectors in this case.

𝓞(*ϵ*^2^) :

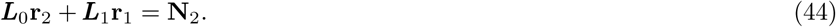

Given the solution *r*_1_, the forcing term has the form

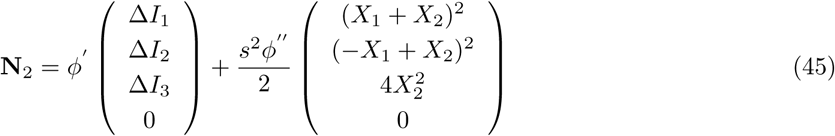

This forcing term contains a projection in the left-null eigenspace of ***L***_0_. This projection cannot give rise to a solution in **r**_2_ and so must be eliminated. The solvability conditions are therefore

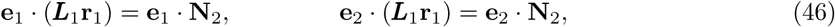

which yield Eqs.32. Note that the resulting normal form has detuning or bias terms, proportional to differences in inputs; these would be called ‘drifts’ in the context of a DDM, but no term linearly proportional to the normal form amplitudes, or ‘decision variables’, as in the normal form for 2-choice DM. Such a term describes the growth rate of the critical mode away from the bifurcation point: attracting to one side and repelling to the other. To derive these terms we will need to continue the calculation to next order. First, we must solve for r_2_. We note that while r_2_ cannot have components in the directions e_1_ or e_2_, components in the directions e_*C*_ = (1, 1, 1, 0) and e_*I*_ = (0, 0, 0, 1) (the common and inhibitory modes which we discussed in the section of DDMs) can be solved for.

Therefore we take **r**_2_ = e_*C*_ *R*_*2C*_ + e_*I*_ *R*_*2I*_ and project

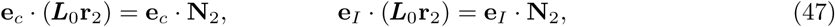

which yields

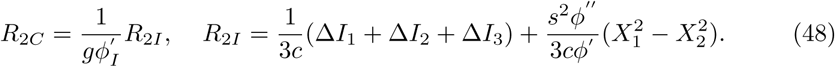

𝓞(*ϵ*^3^) :

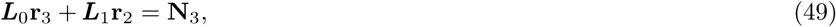

where

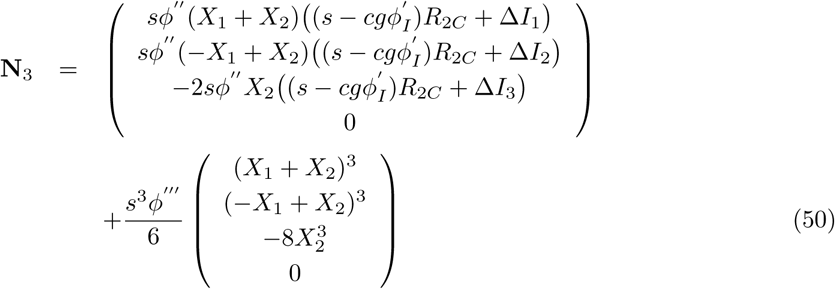

Applying the solvability conditions gives the equations

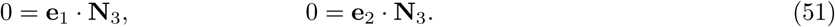

If we add the resulting terms to the normal form equations derived at 𝓞(*ϵ*^2^) we have

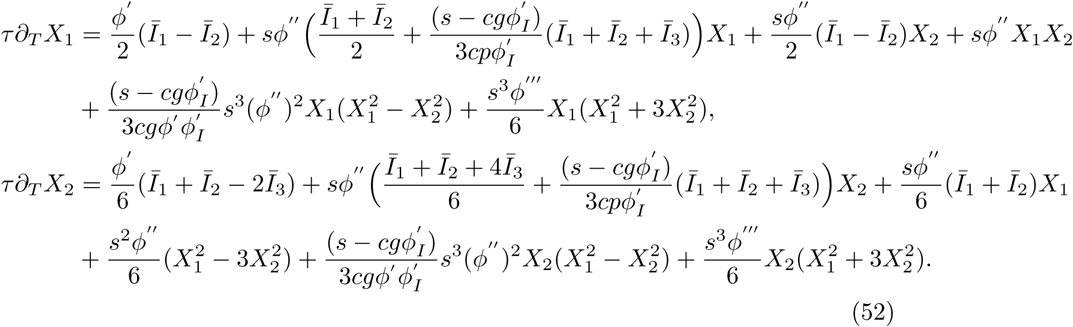

#### Normal form for 3-choice DM starting with a different neuronal model

If we consider a different neuronal model for 3-choice DM we can still arrive at the same normal form equations. The only important factor is the presence of a linear subspace associated with a zero eigenvalue of multiplicity two and which is spanned by eigenvectors representing competition modes. As an example, consider the model

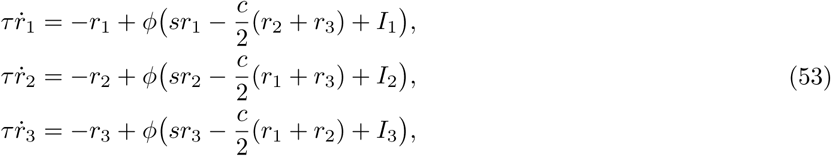

in which the inhibition is not explicitly modelled as before. A linear stability analysis reveals that there is a zero eigenvalue with multiplicity two when 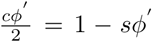 and 1 − *sφ′ >* 0. Therefore, the instability mechanism here is the cross-inhibition, while before it was the self-coupling. Nonetheless, it is only the structure of the corresponding eigenvectors which matters. In this case the two eigenvectors with zero eigenvalue are e_1_ = (1, −1, 0) and e_2_ = (1, 1, −2). Following the methodology described above leads to the identical normal form equations Eqs.32 with *s* replaced by *s* + *c/*2.

### Normal form equations for n-choice DM

A neuronal model for n-choice DM (*n >* 3) is

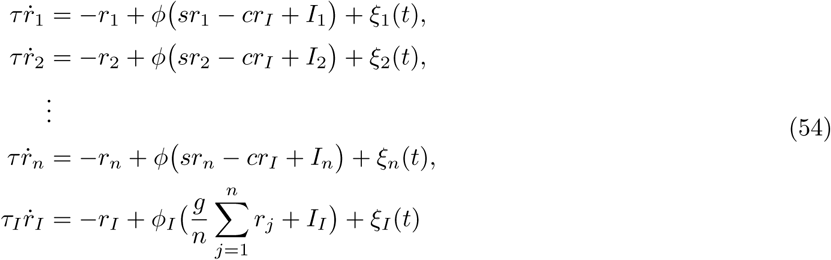

We again consider small differences in the inputs, *I*_i_ = *I*_0_ + *ϵ*^2^∆*I*_i_ for *i* = [1*, n*], expand the rates as **r** = **R**+*ϵ***r**_1_+𝓞(*ϵ*^2^), where **R** are the rates at the fixed point, and define the slow time *T* = *ϵt*. If we take 1 − *sφ′* = 0 then the matrix of the linearized system has a zero eigenvalue with multiplicity *n* − 1. Carrying out an expansion up to 𝓞(*ϵ*^2^) as described above and applying the *n* − 1 solvability conditions leads to the normal form equations Eqs.34.

## Declarations

a) Ethics Approval and Consent to Participate

Not applicable.

b) Consent for publication

Not applicable

c) Availability of data and supporting materials

Data sharing not applicable to this article as no datasets were generated or analysed during the current study.

d) Funding

Grant number MTM2015-71509-C2-1-R from the Spanish Ministry of Economics and Competitiveness. Grant 2014 SGR 1265 4662 for the Emergent Group “Network Dynamics” from the Generalitat de Catalunya. This work was partially supported by the CERCA program of the Generalitat de Catalunya.

## Acknowledgements

I acknowledge helpful conversations with Klaus Wimmer.

## Footnotes

[1] The second derivative of the neuronal transfer function at the bifurcation *φ*” will generically be positive if the neurons operate in the fluctuation-driven regime at low to moderate firing rates. Saturating firing rates, which would lead to *φ*” < 0, only occur at unrealistically high firing rates.

[2] Changing the mean input leads to higher-order corrections which can be calculated straightforwardly, see the Appendix.

## List of Abbreviations

DDM: (Drift Diffusion Model)
DM: (Decision Making)

## References

1. Wald, A.: Sequential tests of statistical hypotheses. Ann. of Math. Stat. 16, 117–186 (1945)

2. Ratcliff, R.: A theory memory retrieval. Psychological Review 85, 59–108 (1978)

3. Ratcliff, R., McKoon, G.: The diffusion decision model: theory and data for two-choice decision tasks. Neural Comput. 20, 873–922 (2008)

4. Shadlen, M.N., Kiani, R.: Decision making as a window on cognition. Neuron 80, 791–806 (2013)

5. Kira, S., Yang, T., Shadlen, M.N.: A neural implementation of wald’s sequential probability test. Neuron 84, 861–873 (2015)

6. Roitman, J.D., Shadlen, M.N.: Response of neurons in the lateral intraparietal area during a combined visual discrimination reaction time task. J. of Neurosci. 22(21), 9475–9489 (2002)

7. Wang, X.-J.: Pacemaker neurons for the theta rhythm and their synchronization in the septohippocampal reciprocal loop. J. Neurophysiol. 87, 889–900 (2002)

8. Wong, K.F., Huk, A.C., Shadlen, M.N., Wang, X.-J.: Neural circuit dynamics underlying accumulation of time-varying evidence during perceptual decision-making. Frontiers in Computational Neuroscience (in press) (2007)

9. Churchland, A.K., Kiani, R., Shadlen, M.N.: Decision-making with multiple alternatives. Nature Neuroscience 11, 693–702 (2008)

10. Bogacz, R., Brown, E., Moehlis, J., Holmes, P., Cohen, J.D.: The physics of optimal decision making: a formal analysis of models of performance in two-alternative forced-choice tasks. Psychological Review 113(4), 700–765 (2006)

11. Roxin, A., Ledberg, A.: Neurobiological models of two-choice decision making can be reduced to a one-dimensional nonlinear diffusion equation. PLoS Comp. Biol. 4, 1000046 (2008)

12. McMillen, T., Holmes, P.: The dynamics of choice among multiple alternatives. J. Math. Psychol. 50, 30–57 (2006)

13. Usher, M., Olami, Z., McClelland, J.L.: Hick’s law in a stochastic race model with speed-accuracy tradeoff. J. Math. Psychol. 46, 704–715 (2002)

14. Shadlen, M.N., Newsome, W.T.: Neural basis of a perceptual decision in the parietal cortex (area lip) of the rhesus monkey. J. Neurophysiology 86, 1916–1936 (2001)

15. Wiggins, S.: Introduction to Applied Nonlinear Dynamical Systems and Chaos. Springer, ??? (2003)

16. Ueltzhoeffer, K., Armbruster-Genç, D.J.N., Flebach, C.J.: Stochastic dynamics underlying cognitive stability and flexibility. PLoS Comp. Biol. 11, 1004331 (2015)

17. Nguyen, K.P., Josić, K., Kilpatrick, Z.P.: Optimizing sequential decision in the drift-diffusion model. J. Math. Psychol. 88, 32–47 (2019)

